# Growth in early infancy drives optimal brain functional connectivity which predicts cognitive flexibility in later childhood

**DOI:** 10.1101/2024.01.02.573930

**Authors:** Chiara Bulgarelli, Anna Blasi, Samantha McCann, Bosiljka Milosavljevic, Giulia Ghillia, Ebrima Mbye, Ebou Touray, Tijan Fadera, Lena Acolatse, Sophie E. Moore, Sarah Lloyd-Fox, Clare E. Elwell, Adam T. Eggebrecht, BRIGHT Study Team

**Author notes:** The BRIGHT team are (in alphabetic order): Muhammed Ceesay, Kassa Kora, Fabakary Njai, Andrew Prentice, Mariama Saidykhan. **Corresponding Author:** Dr. Chiara Bulgarelli, Address: Centre for Brain and Cognitive Development Department of Psychological Sciences, Birkbeck, University of London Malet Street, London, WC1E 7HX (UK).

## Abstract

Functional brain network organization, measured by functional connectivity (FC), reflects key neurodevelopmental processes for healthy development. Early exposure to adversity, e.g. undernutrition, affects neurodevelopment, observable via disrupted FC, and leads to poorer outcomes from preschool age onward. We assessed longitudinally the impact of early growth trajectories on developmental FC in a rural Gambian population from age 5 to 24 months. To investigate how these early trajectories relate to later childhood outcomes, we assessed cognitive flexibility at 3-5 years. We observed that early physical growth before the fifth month of life drove optimal developmental trajectories of FC that in turn predicted cognitive flexibility at pre-school age. In contrast to previously studied developmental populations, this Gambian sample exhibited long-range interhemispheric FC that decreased with age. Our results highlight the measurable effects that poor growth in early infancy has on brain development and the possible subsequent impact on pre-school age cognitive development, underscoring the need for early life interventions throughout global settings of adversity.

## Introduction

The first 1000 days of life are of paramount importance for human brain (1) and body development (2, 3) Early adversity negatively impacts infant development, which leads to long-term consequences from delayed milestones in childhood development to lowered productivity at the individual and societal levels (4–6). Therefore, assessing trajectories of brain development in infants exposed to early adversity is essential to understand and mitigate effects on brain development and subsequent outcomes at school age and beyond (7, 8).

Early undernutrition engenders detrimental effects on a range of cognitive skills (9), with long-lasting effects through adulthood (10). Interventions using therapeutic feeding regimens and supplements have been shown to have a strong causal impact on childhood undernutrition, brain development, and subsequent neurodevelopmental outcomes (11–13). Despite numerous attempts of interventions, global rates of undernutrition remain high (14), with infants in low- and middle-income countries (LMICs) at greatest risk (15). Up to one-third of infants in LMICs are at risk of not meeting standard milestones of social, motor, linguistic, and cognitive development by pre-school age (16, 17), which can lead to lifelong consequences that impact social and economic development at individual, national, and global scales (18). Thus, it is essential to study neurodevelopment throughout the first two years in regions with high rates of undernutrition in order to optimize early interventions. To identify effective early interventions, we must first identify the mechanisms that link undernutrition to later cognitive outcomes. Effects of undernutrition on the developing brain may manifest through alterations in function brain networks (19), as shown in school-age children exposed to chronic poverty (20). However, our understanding of the relationships between early undernutrition, brain connectivity, and later cognitive outcomes is still very limited.

Functional brain network organization is commonly assessed with resting-state functional connectivity (FC) via temporal correlations in brain activity as measured with functional magnetic resonance imaging (fMRI) (21, 22). Based on data from fMRI, current models hypothesize that FC patterns mature throughout early development (23–27), where in typically developing brains, adult-like networks emerge over the first years of life as long-range functional connections between pre-frontal, parietal, temporal, and occipital regions become stronger and more selective (28–31). This maturation in FC has been shown to be related to the cascading maturation of myelination and synaptogenesis (32, 33) - fundamental processes for healthy brain development (34). Therefore, disrupted patterns of FC may reflect disrupted anatomical maturation of brain circuits and systems. For example, preterm infants with severe diffuse white matter impairment exhibited lower levels of FC in executive function networks compared with preterm infants with no or mild white matter impairment (35). Importantly, normative developmental patterns may be disrupted and even reversed in clinical conditions that impact development; e.g., increased short-range and reduced long-range FC have been observed in preterm infants (36) and in children with autism spectrum disorder (37, 38).

While widely used neuroimaging modalities such as fMRI can elucidate typical and atypical developmental trajectories, they are often poorly suited for lower resource neuroimaging contexts due to limited portability, need for specialized facilities, and the high cost associated with these methods (28, 36, 39). To address the need for neuroimaging studies in lower resource environments including (but not limited to) in LMICs, researchers have increasingly turned to more portable tools, including electroencephalography (EEG) (40), functional near-infrared spectroscopy (fNIRS) (41–43) and diffuse correlation spectroscopy (DCS) (44). These tools enable the creation of mobile neuroimaging laboratories that can be deployed virtually anywhere, eliminating practical constraints imposed by costly and immobile methods. Indeed, research based in The Gambia, Guinea-Bissau, Bangladesh and Ivory Coast have established feasibility of these tools to perform brain imaging studies outside of a specialized lab environment, with recent studies reporting altered cortical physiology related to early adversity and undernutrition (40, 43–45). For example, using EEG, Xie and colleagues showed that in 2-3 year old Bangladeshi children, growth faltering was associated with FC in the theta and beta frequency bands, which was negatively related to children’s IQ score at 4 years (46). With regards to fNIRS, it has a better spatial resolution and anatomical specificity than EEG, thus providing more precise and reliable localization of brain networks (47, 48). Moreover, fNIRS facilitates testing in awake and engaged infants, making the results more comparable with adult FC studies that are performed on awake participants (49). Given these unique strengths, fNIRS has been used to study FC in longitudinal studies in awake infants in high-resource settings (31), and the portability and the low cost of this equipment has fostered implementation outside conventional labs and in LMICs (50). Because fNIRS overcomes logistical (MRI) and resolution (EEG) challenges of other common modalities, we chose it for this study to assess early life brain health of children in LMIC exposed to early severe undernutrition.

The goal of the study was to investigate early physical growth in infancy, developmental trajectories of brain FC across the first two years of life, and cognitive outcome at school age in a longitudinal cohort of infants and children from rural Gambia, an environment with high rates of maternal and child undernutrition. Specifically, we aimed to(i)investigate whether differences in physical growth through the first two years of life are related to FC at 24 months, and (ii) investigate if early FC has an impact on cognitive outcome at pre-school age in these children. Data were collected from N=204 children as part of the Brain Imaging for Global Health project (BRIGHT; globalfnirs.org/the-bright-project), a longitudinal study examining infant development from term birth to 24-months of age. We acquired longitudinal measurements of brain FC in awake Gambian infant participants at five age points (5, 8, 12, 18, and 24 months) using fNIRS as the infants watched calming videos. We also acquired anthropometric measures (i.e., weight and length) as indicators of nutritional status at birth, at 7-to-14 days, one month of age, and at all five fNIRS data acquisition time points. Additionally, because developmental delays interfere with everyday functioning starting within the pre-school age range (16), we assessed cognitive flexibility, a core executive function (51), at ages three and five years. We hypothesized that (i) long-range FC would increase and short-range FC would decrease throughout the first two years of life, (ii) that early positive physical growth would significantly predict long-range FC, and that (iii) long-range FC strength would predict cognitive flexibility performance in pre-schoolers. Our results provide novel insights into the developmental trajectory of the functional organization of the developing brain, its relationship with nutritional adversity exposure and status, and the subsequent effects on preschool-age cognitive outcomes.

## Results

### Gambian infants exhibit robust FC development throughout first two years

The repeated assessments of brain FC throughout the first two years of life allowed us to analyse developmental trajectories of the maturation of early brain networks and how they relate to measures of infants’ growth as well as to cognitive flexibility at preschool age (**Figure 1A**). To estimate how FC changed between 5-24 months of age in Gambian children, we performed linear mixed models (LMM) using data from the 132 infants with valid data for at least two time points. To increase statistical power while minimizing errors due to variability in head anatomy and cap placement, we divided the array into six sections and calculated functional connectivity (FC) between all possible 21 interhemispheric homotopic, intrahemispheric within section, fronto-posterior, and crossed connections (**Figure SI1**). Results on the Fisher-z transformed correlation coefficients (z-RHO) of the oxygenated haemoglobin (HbO_2_) showed that frontal interhemispheric homotopic FC decreased with age (*F*=11.03, *p*<0.001), and that left (*F*=5.8, *p*<0.001) and right fronto-middle (*F*=4.86, *p*<0.001) and right fronto-posterior (*F*=5.52, *p*<0.001) FC increased with age (**Figure 2**).

**Figure 1.**
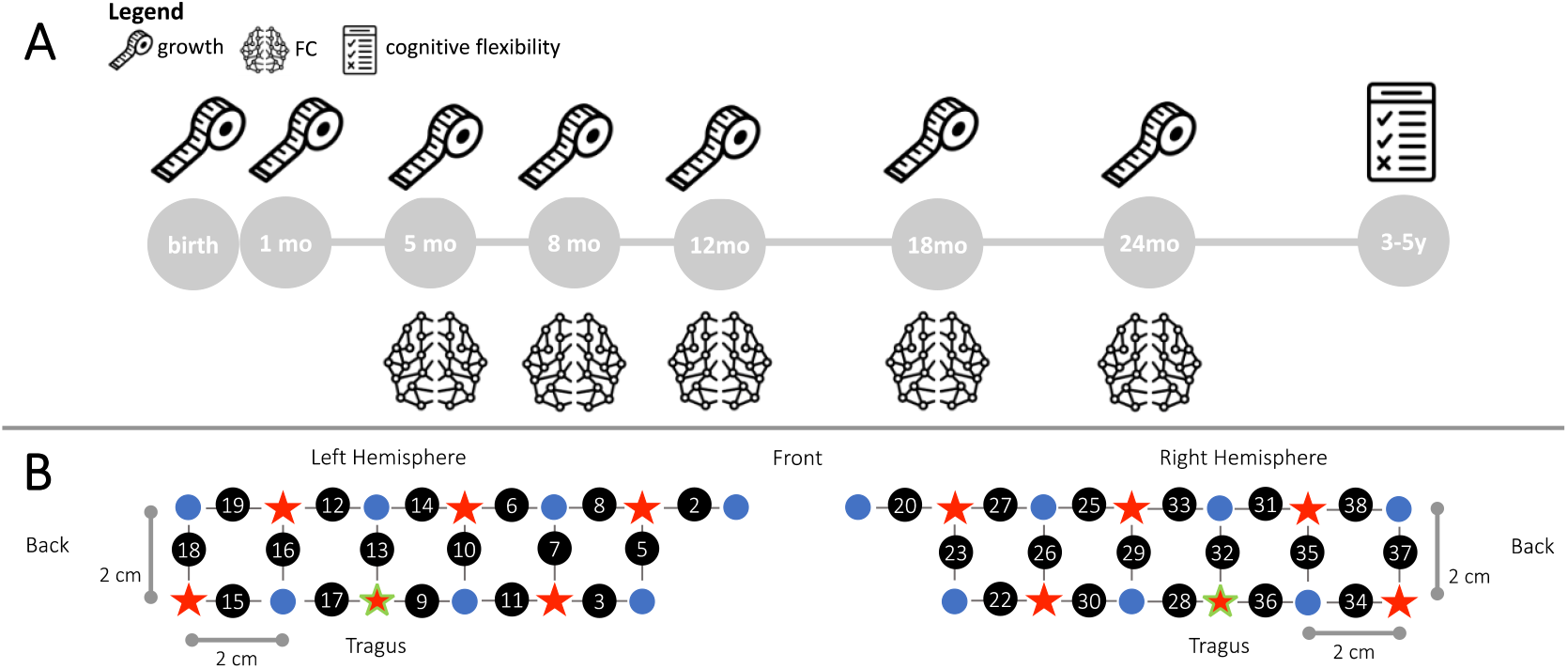
Experimental design. (A) Measures taken in the BRIGHT project used in this work. The measuring tape represents anthropometric measures, the brain represents fNIRS FC and the test represents the cognitive flexibility assessment. (B) Schematic representation of the spatial layout of the fNIRS array. Sources are marked with red stars, detectors are marked with blue circles, channels are marked with grey lines and numbered with black circles. The channels/optodes used as a reference for the tragus are highlighted in green.

**Figure 2.**
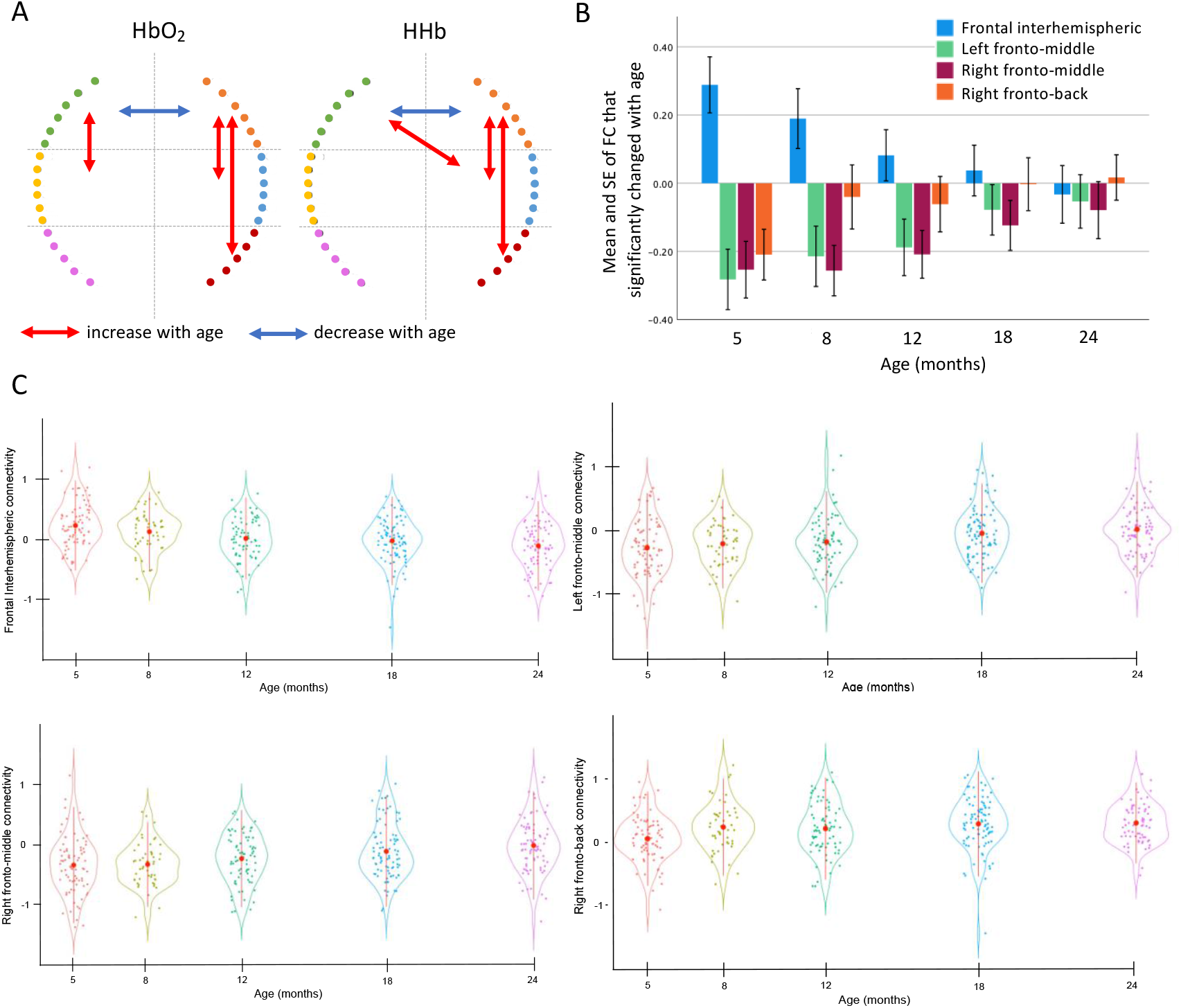
Linear mixed models results showing FC that displayed a statistically significant change with age. (**A**) Significant results of the linear mixed model, blue indicates connections that decreased with age, red indicates connections that increase with age. (**B**) Mean and standard error of the mean (SE) of the functional connections that changed with age (HbO_2_). Error bars are 1 SE. (**C**) Violin plot showing the mean ± standard deviation (SD) (red circles and lines) and the individual variability (coloured dots) of the FC that showed a change with time (HbO_2_).

Results on the z-RHO scores of the deoxygenated haemoglobin (HHb) largely agreed with the results on HbO_2_ (**Table 1 and Figure 2**). Frontal interhemispheric FC significantly decreased with age, showing positive correlations at five months, and negative values at 24 months (**Figure 2B and Figure 2C**). The FC between fronto-middle and frontal-posterior that significantly increased with age showed strong anticorrelation values at early ages that gradually decreased but were still negative at 24 months (**Figure 2B and Figure 2C**).

**Table 1.**
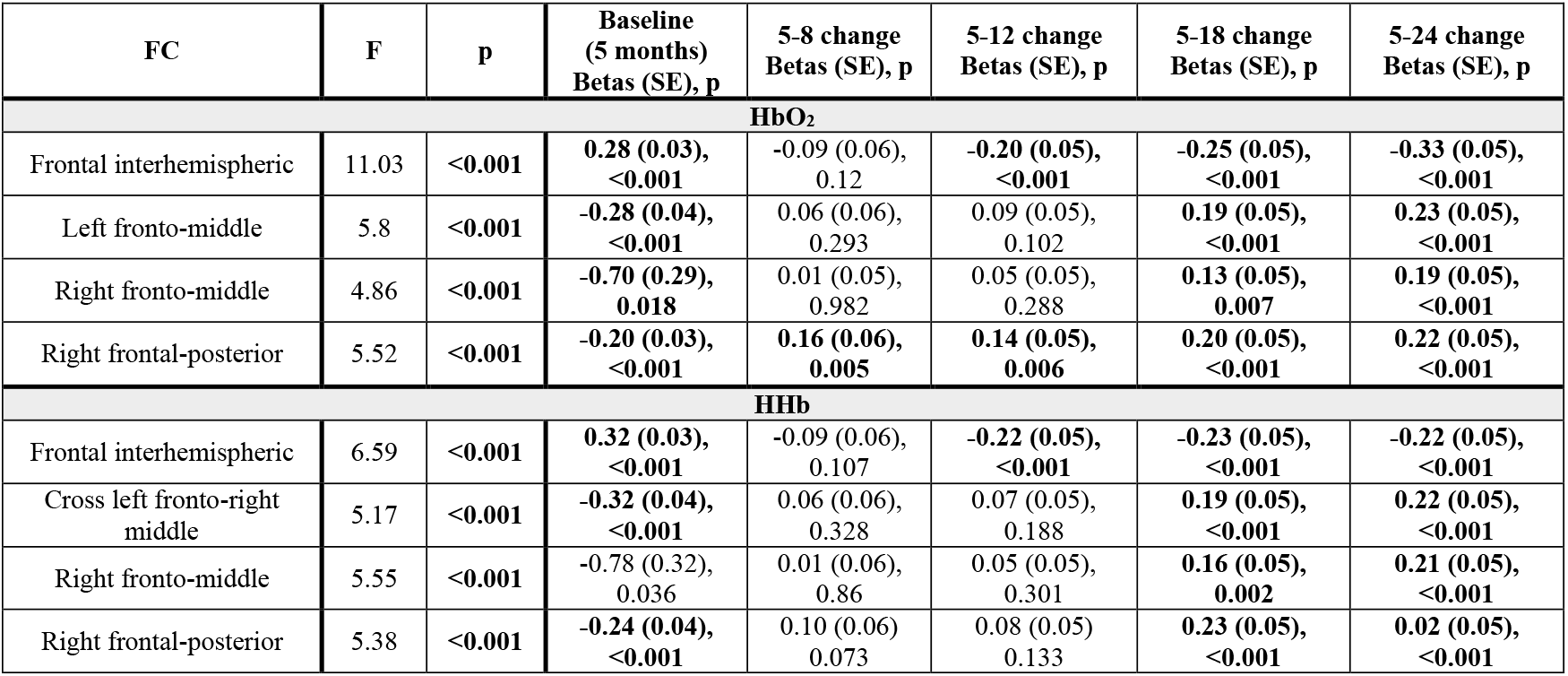
FC that significantly changed with age. Results are displayed in terms of estimated betas, standard errors and p values that survived Bonferroni correction. Regressions that survived Bonferroni correction for multiple comparisons are in bold.

Notably, results from LMMs performed on data pre-processed without the global signal regression (GSR) (see results in SI, **Table SI1** and **Figure SI2**) confirmed a decreasing trajectory of the frontal interhemispheric homotopic FC and an increasing trajectory of bilateral short-range FC with age.

### Early growth trajectories predict functional connectivity at 24 months

To investigate the impact of early nutritional status on FC at 24 months, we used multiple regression with the infant growth trajectory (delta weight for length z-score between all time points, ΔWLZ) and FC at 24 months, adjusting for WLZ at birth or head-circumference z-score (HCZ) at 7/14 days. To maximise power, we considered only those FC that showed a statistically significant change with age. The ΔWLZ between birth/1 month and older ages positively predicted frontal interhemispheric homotopic FC at 24 months (ΔWLZ birth with older ages all *p*<0.02 FDR corrected, ΔWLZ 1 month with older ages all *p*<0.03, see **Figure SI3** for some examples of scatterplots) and negatively predicted left and right fronto-middle FC at 24 months (*p*<0.04). The ΔWLZ between birth and 1 month positively predicted right fronto-posterior FC at 24 months (*p*=0.006, FDR corrected) while ΔWLZ between 1 month and older ages negatively predicted right fronto-posterior FC at 24 months (all *p*<0.01, FDR corrected). Interestingly, ΔWLZ between 5/8/12/18 months and older ages did not show statistically significant impacts on any of the FC assessed (**Table 2 and Figure 3A**).

**Table 2.** Results of the regression analyses of the effect of ΔWLZ on FC at 24 months. Significant positive associations are in green, significant negative associations are in orange, and non-significant (NS) associations are in blue; * indicates regressions that are still significant after correcting for HCAZ at 7/14 days and ** indicates regressions that are still significant after correcting for neonatal HCAZ and WLZ. Regressions that survived FDR correction for multiple comparisons are in bold.

| Frontal interhemispheric FC at 24 mo |  |  |  |  |  |  |  |
| --- | --- | --- | --- | --- | --- | --- | --- |
| ΔWLZ | birth | 1 mo | 5 mo | 8 mo | 12 mo | 18 mo | 24 mo |
| birth | | NS | $F(1,68)=8.69$,<br>$p=0.004^*$,<br>$R^2=0.102$ | $F(1,67)=7.65$,<br>$p=0.007^*$,<br>$R^2=0.104$ | $F(1,69)=10.4$,<br>$p=0.002^{**}$,<br>$R^2=0.121$ | $F(1,68)=10.98$,<br>$p=0.001^{**}$,<br>$R^2=0.142$ | $F(1,59)=5.40$,<br>$p=0.024^*$,<br>$R^2=0.085$ |
| 1 mo | | | $F(1,77)=5.59$,<br>$p=0.021^{**}$,<br>$R^2=0.069$ | NS | $F(1,78)=4.74$,<br>$p=0.032^{**}$,<br>$R^2=0.058$ | $F(1,75)=4.45$,<br>$p=0.038^{**}$,<br>$R^2=0.057$ | NS |
| 5 mo |  |  |  | NS | NS | NS | NS |
| 8 mo |  |  |  |  | NS | NS | NS |
| 12 mo |  |  |  |  |  | NS | NS |
| 18 mo |  |  |  |  |  |  | NS |
| 24 mo |  |  |  |  |  |  |  |
| Left fronto-middle FC at 24 mo |  |  |  |  |  |  |  |
| ΔWLZ | birth | 1 mo | 5 mo | 8 mo | 12 mo | 18 mo | 24 mo |
| birth | | NS | NS | $F(1,67)=4.07$,<br>$p=0.048^{**}$,<br>$R^2=0.058$ | NS | NS | $F(1,59)=4.24$,<br>$p=0.044^{**}$,<br>$R^2=0.068$ |
| 1 mo |  |  | NS | NS | NS | NS | NS |
| 5 mo |  |  |  | NS | NS | NS | NS |
| 8 mo |  |  |  |  | NS | NS | NS |
| 12 mo |  |  |  |  |  | NS | NS |
| 18 mo |  |  |  |  |  |  | NS |
| 24 mo |  |  |  |  |  |  |  |
| Right fronto-middle FC at 24 mo |  |  |  |  |  |  |  |
| ΔWLZ | birth | 1 mo | 5 mo | 8 mo | 12 mo | 18 mo | 24 mo |
| birth | | NS | NS | $F(1,67)=4.74$,<br>$p=0.033$,<br>$R^2=0.067$ | NS | $F(1,68)=5.29$,<br>$p=0.024^{**}$,<br>$R^2=0.073$ | NS |
| 1 mo | | | $F(1,77)=4.26$,<br>$p=0.042^{**}$,<br>$R^2=0.053$ | $F(1,75)=6.66$,<br>$p=0.012^{**}$,<br>$R^2=0.083$ | NS | $F(1,75)=5.33$,<br>$p=0.024^{**}$,<br>$R^2=0.067$ | NS |
| 5 mo |  |  |  | NS | NS | NS | NS |
| 8 mo |  |  |  |  | NS | NS | NS |
| 12 mo |  |  |  |  |  | NS | NS |
| 18 mo |  |  |  |  |  |  | NS |
| 24 mo |  |  |  |  |  |  |  |
| Right frontal-posterior FC at 24 mo |  |  |  |  |  |  |  |
| ΔWLZ | birth | 1 mo | 5 mo | 8 mo | 12 mo | 18 mo | 24 mo |
| birth | | $F(1,66)=8.26$,<br>$p=0.006^{**}$,<br>$R^2=0.113$ | NS | NS | NS | NS | NS |
| 1 mo | | | $F(1,69)=4.86$,<br>$p=0.031^*$,<br>$R^2=0.067$ | $F(1,68)=6.98$,<br>$p=0.01^{**}$,<br>$R^2=0.094$ | $F(1,71)=8.86$,<br>$p=0.004^{**}$,<br>$R^2=0.112$ | $F(1,67)=7.72$,<br>$p=0.007^{**}$,<br>$R^2=0.105$ | $F(1,57)=7.03$,<br>$p=0.01^{**}$,<br>$R^2=0.112$ |
| 5 mo |  |  |  | NS | NS | NS | NS |
| 8 mo |  |  |  |  | NS | NS | NS |
| 12 mo |  |  |  |  |  | NS | NS |
| 18 mo |  |  |  |  |  |  | NS |
| 24 mo |  |  |  |  |  |  |  |

**Figure 3.**
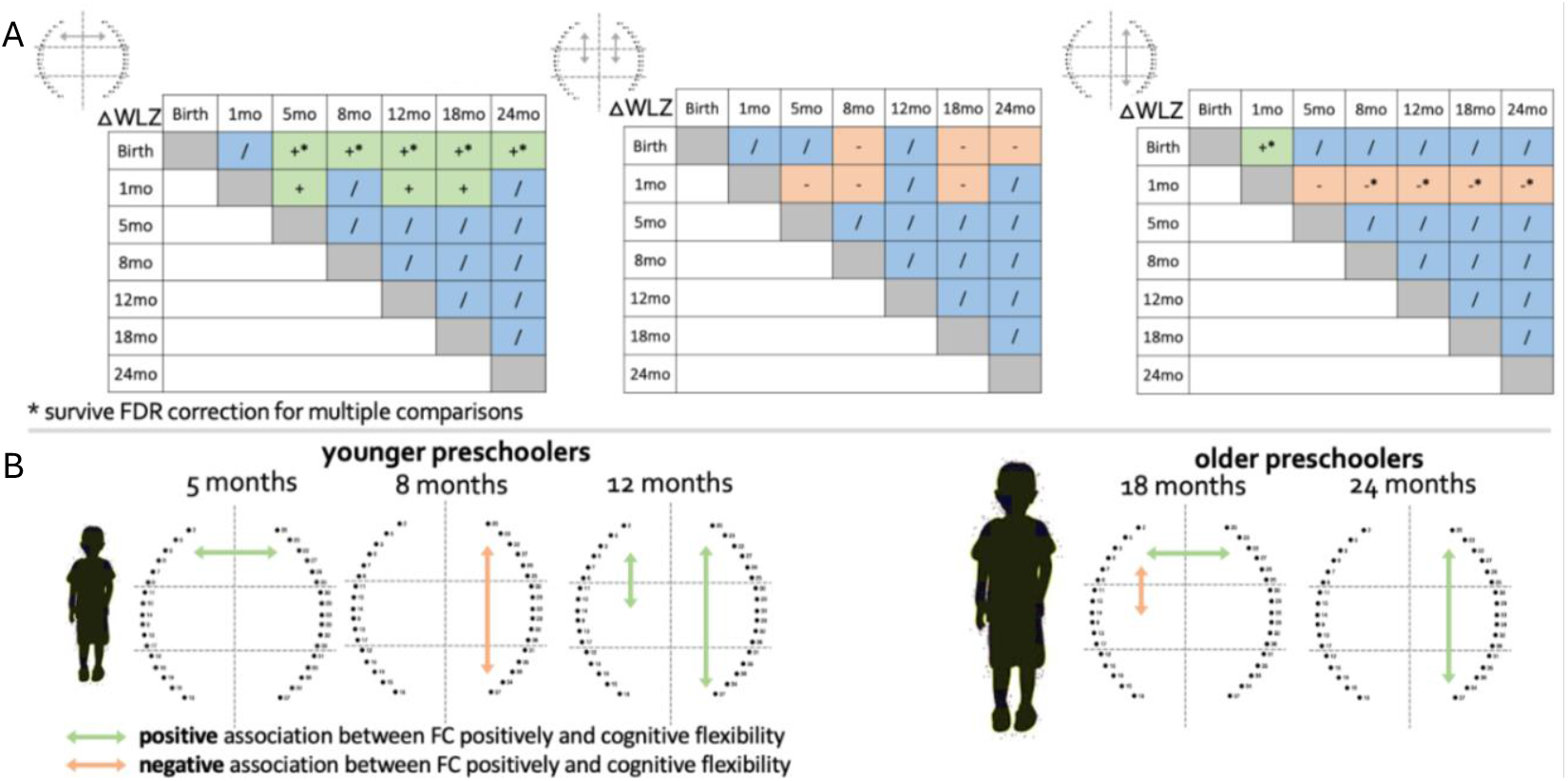
Associations between FC, early growth and later cognitive flexibility. (**A**) Significant positive associations are in green, significant negative associations are in orange, and non-significant associations are in blue. * indicates regressions that survived FDR correction for multiple comparisons. (**B**) Schematic representations of the early FC connections shown to predict cognitive flexibility in preschoolers. Significant positive associations are in green, significant negative associations are in orange (these results did not survive corrections for multiple comparisons).

### Early functional connectivity predicts cognitive flexibility at preschool age

To investigate whether FC early in life predicted cognitive flexibility at preschool age, we used multiple regression of FC across the first two years of life against later cognitive flexibility in preschoolers at three and five years. As per the analysis above, we focused on only those FC that showed a statistically significant change with age. Our results showed that frontal interhemispheric homotopic FC at 5 months (*F*(1,38)=4.21, *p*=0.047, R^2^=0.102), left frontal-middle (*F*(1,33)=7.86, *p*=0.009, R^2^=0.197) and right frontal-posterior FC at 12 months (*F*(1,38)=4.82, *p*=0.034, R^2^=0.115) positively predicted cognitive flexibility in young preschoolers. Right frontal-posterior FC at 8 months negatively predicted cognitive flexibility in young preschoolers (*F*(1,32)=5.6, *p*=0.024, R^2^=0.153). Frontal interhemispheric homotopic FC at 18 months (*F*(1,45)=4.82, *p*=0.030, R^2^=0.115), and right frontal-posterior FC at 24 months (*F*(1,49)=4.85, *p*=0.032, R^2^=0.092) positively predicted cognitive flexibility in older preschoolers. Left front-middle FC at 18 months negatively predicted cognitive flexibility in older preschoolers (*F*(1,45)=5.72, *p*=0.021, R^2^=0.115). While these associations between early functional connectivity and cognitive flexibility at preschool age show promise, none of these survived FDR correction for multiple comparisons (**Figure 3B and Table SI2**), and so should be considered as preliminary and useful for hypothesis generation for future studies.

We also explored whether changes in growth and changes in functional connectivity between 5 and 24 months were associated with cognitive flexibility at preschool age, but we did not find any significant association (**Table SI3** and **Table SI4**).

## Discussion

This study investigated how early adversity via undernutrition drives brain functional connectivity throughout the first two years of life and how these early functional connections are associated with cognitive flexibility at preschool age. To capture the rapid neural development that takes place during early years in community LMIC settings we used fNIRS to assess brain connectivity (22, 24, 26, 31). Additionally, we recorded brain activity while infants were awake, thus making the results more comparable with those from adult studies. Our results show: (i) healthy growth trajectories during early infancy are crucial for healthy FC at 24 months; (ii) our cohort of Gambian infants exhibit atypical developmental trajectories of functional connectivity (FC) in over the first two years of life; and (iii) FC during the first two years of life may predict cognitive flexibility at preschool age.

First, we found that while left and right intrahemispheric fronto-middle and right frontal-posterior FC increased with age, frontal inter-hemispheric FC decreased with age. Additionally, we observed long-range right frontal-posterior FC increased with age and exhibited positive correlations at 24 months, as consistent with previous infant studies (29, 31). Previous findings from typically developing infants measured in the US showed that long-range FC gradually strengthened with age, indicating maturing of functional networks that span distant brain regions (28, 29). Moreover, the presence of inter-hemispheric connections are known to indicate healthy development of FC (52) consistent with the continuous development of the corpus callosum from infancy until early adulthood (53).

Conversely, decreased inter-hemispheric FC has been observed where development can follow a more divergent and/or heterogeneous path, such as in autism (54). The observed decrease in frontal inter-hemispheric FC with increasing age may be due to the exposure to early life undernutrition adversity. We acknowledge that differences in FC could also be attributed to other environmental and cultural disparities between high-resource and low-resource settings, and future studies are needed to explore this further. Finally, our results indicated that the Gambian infant bilateral fronto-middle FC exhibited anticorrelation at five months, and then gradually became less negative with age. This pattern suggests that this fronto-middle FC may become in sync later after the second year of life. The fNIRS array covers regions over the front and middle cortical regions, putatively belonging to the default mode network and fronto-parietal network, which are normally anticorrelated (55), suggesting that the earlier observed anticorrelation pattern might indeed be adaptive.

Importantly, these results remained significant even without GSR, indicating that our findings are not solely driven by preprocessing choices. While the use of GSR in FC studies remains debated (56), in the absence of short channels (which are difficult to use reliably with infants (57)) and external physiological measures, applying GSR represented the most appropriate preprocessing option. In fact, failure to correct for systemic physiological fluctuations can, in fact, lead to artificially elevated connectivity estimates in fNIRS data (58). While further investigations are needed to confirm these findings, these results highlight a general disruption of the integration/segregation process of network development among the infants in our study (59).

Second, a central goal of the current work was to investigate whether infant nutritional status, as measured by change in weight for length z-scores (ΔWLZ), was associated with functional connectivity at 24 months. First, we observed that ΔWLZ between neonatal time points and older ages positively predicted frontal interhemispheric homotopic FC at 24 months, which supports the idea that the decreasing interhemispheric connectivity observed with age might be atypical, as greater increases in ΔWLZ were found to be positively associated with stronger interhemispheric FC. This is consistent with research showing an impact of nutritional status on long-range connections and brain networks development (60–62) and the decreased long-range FC found in preterm babies with faltering growth (63–65). A common feature of the observed associations between ΔWLZ and FC is that they were statistically significant only when ΔWLZ was calculated including measures collected neonatally (i.e., at birth or one month of age). Changes in growth later in development (i.e. ΔWLZ between 5, 8, 12, or 18 months and older ages) did not show statistically significant impacts on any of the FC assessed. This provides support for the hypothesis that undernutrition during the first months of life is more impactful on brain development than undernutrition occurring later in infancy and early childhood. While the impact of undernutrition on brain development has been documented in LMICs (46), herein, we provided empirical evidence that growth faltering specifically in infants younger than five months of age impacts observable development of functional brain networks in the second year of life. Future studies may be needed to pinpoint which specific brain networks are impacted. This result also suggests that early undernutrition has a lasting impact on subsequent cognitive skills, even in children who show subsequent catch-up growth.

The other primary aim of this study was to investigate whether early FC development was related to cognitive flexibility in preschoolers at three and five years of age. Long range FC (interhemispheric homotopic and frontal-posterior) positively predicted performance in the cognitive flexibility task both in younger and older preschoolers. This is consistent with the fact that the integration of distant brain regions fosters cognitive development, especially in prefrontal cortex (66, 67), possibly driven by maturation within the fronto-parietal network that has been widely documented as a primary neural underpinning of cognitive flexibility (68). Additionally, left fronto-middle FC at 12 months positively predicted cognitive flexibility in young preschoolers, but the same connection at 18 months negatively predicted cognitive flexibility in older preschoolers. This shift in direction of the relationship between FC and cognitive flexibility might indicate the shift between short-range and long-range FC during development, where short-range FC still promotes cognitive development early in life, while becoming less impactful as the child ages possibly because children develop a wider range of strategies to support cognitive flexibility as they age. A recent review on the neural underpinnings of cognitive flexibility highlighted that right-lateralized activations underlying this skill increased with age, while activations over the left hemisphere decreased with age (51). Our findings are consistent with this view, suggesting an increase in functional specialisation and segregation with development. While our results are consistent with previous studies, we acknowledge that the significant associations between early FC and later cognitive flexibility do not withstand multiple comparisons. Therefore, we encourage future studies that may replicate these findings with a larger sample.

Some limitations of the current study merit discussion. Although the BRIGHT sample is one of the largest longitudinal neuroimaging studies in LMIC to assess longitudinally this many age points during the first two years of life, the sample size may still be underpowered to assess the modest effects of the regression models. Second, additional factors may affect FC that were not considered in our models. Applying more advanced statistical modelling methods and structural equation modelling analyses may provide greater insight with further investigations in contexts of adversity and, in turn, establish which outcomes are predicted by FC. That said, results from the regression models were statistically robust, as most survived FDR correction for multiple comparisons, and were significant after including HCZ at 7-14 days and WLZ at birth in the models. It is also worth noting that in line with previous FC fNIRS studies (31, 69), linear mixed model results on the HbO_2_ and the HHb signals showed consistency in the FC changes with age measured in the two chromophores, and similar trajectories when removing the global signal regression step from the pre-processing, suggesting that the results were reliable and statistically robust. Future collaborations between projects studying the impact of adversity on brain development in both high resource and LMIC settings are crucial to advance the field and therefore inform efficient strategies – including the timing - of intervention. Even given the large sample in our study, we were underpowered to test for group comparisons between sets of infants with distinct undernutrition growth profiles, e.g., infants with early poor growth that later resolved and infants with standard growth early that had a poor growth later. We were also underpowered to test the associations between early growth and FC on clinically undernourished infants (defined as having ΔWLZ two standard deviations below the mean). However, our findings provide a solid foundation for future research on the impacts of undernutrition during the first months of life on later brain health and cognitive abilities throughout childhood.

In conclusion, herein we showed that, as expected, positive growth trajectories during the first few months of life were positively associated with stronger interhemispheric frontal FC and negatively associated with bilateral short-range FC. Interestingly, while Gambian infants had expected increasing left and right intrahemispheric fronto-middle and right frontal-posterior FC with age, frontal inter-hemispheric FC decreased with age, which is inconsistent with extant data from other populations studied in Europe and US. Of particular note, growth during the first few months after birth showed a significant impact on brain health, as measured with FC, while later growth changes were not related to variability in FC. Moreover, long-range FC predicted the performance on a cognitive flexibility task at preschool age. Taken together, our results show the impact that early growth has on functional brain development, which in turn affects cognitive flexibility in preschoolers.

Finally, this study emphasizes the importance of the timing of interventions, for which benefits are visible at preschool age and beyond.

## Materials and Methods

### Experimental Design

Our study investigated the developmental trajectory of brain health as assessed with functional connectivity (FC) in Gambian infants and its relationship with early growth, as a proxy for nutritional status, and later cognitive outcomes. To do, so we estimated FC from data acquired with fNIRS longitudinally at five time points between age five and 24 months in the context of the BRIGHT Project. The BRIGHT project sought to provide brain function-for-age curves from the UK and Gambian infants and to gain insight into the effects that issues related to living in low-resource settings may have on brain development. We then tested for relationships between brain FC at age 24 months with measures of early growth, as indexed by changes in weight-for-length z-scores (reflecting body weight in proportion to attained growth in length) at one month of age, and at each of the four subsequent visits (details provided below). We also tested for associations between brain FC at each time point with later cognitive outcome, as indexed by cognitive flexibility assessed in three- and five-year-olds preschool-age children (**Figure 1A**). Ethical approval was granted by the joint Gambia Government - MRC Unit the Gambia Ethics Committee, and written informed consent was obtained from all parents prior to participation.

### Participants

We used data from N=204 infants recruited in rural community settings of The Gambia as part of the BRIGHT project. The study was a prospective, longitudinal study with fNIRS assessments planned for awake infants at ages 5, 8, 12, 18 and 24 months. All infants were born full term (37–42 weeks gestation). fNIRS data were additionally acquired from 1-month-old infants while asleep. However, as sleep stages may impact connectivity measures (98), these data are not part of the current analysis. Reasons for exclusion from the fNIRS analysis were: i) fussiness; ii) experimental errors (missing pictures of the headgear placement or any other technical issue); iii) poor headgear placement (see supplementary information for *Assessment of headgear placement*); iv) low-quality data (more than 40% of the channels excluded as previously done (70–73)). **Table SI5** summarises the details of the included and excluded participants for each visit, showing an attrition rate within the standard range for infant fNIRS studies (74) (see **Table 3** for the demographic information of the participants included in the analyses at each age).

**Table 3.** Demographic information per each age.

| **Age** | **Age in days (mean±SD)** | **Sex (M, F)** | **WLZ (mean±SD)** | **HCZ (mean±SD)** |
| --- | --- | --- | --- | --- |
| **5 months** | 158.92±9.91 | 45,42 | -0.24±0.95 | -0.72± 0.92 |
| **8 months** | 249.34±17.97 | 28,25 | -0.24±1.01 | -0.81±0.90 |
| **12 months** | 372.92±14.75 | 42,40 | -0.58±1.05 | -0.95±0.95 |
| **18 months** | 560.37±24.38 | 48,49 | -0.85±0.91 | -0.82±0.93 |
| **24 months** | 746.61±23.25 | 49,47 | -0.44±0.99 | -0.80±0.97 |

### Anthropometric measures

Measurements of length, weight, and head circumference (HC) were collected on all infants at birth, at 7/14 days, 1 month, and in all visits from 5 to 24 months, by trained field workers using calibrated tools. Length was measured using a Harpenden Infantometer length board (Holtain Ltd) to a precision of 0.1 cm. Weight was obtained using an electronic baby scale (Model 336, SECA) to a precision of 0.01 kg. Finally, HC was measured around the maximum circumference of the head (forehead to occiput) using stretch-proof measuring tape (Model 201, SECA) to the nearest 0.1 cm. Each measure was taken in triplicate and the mean of the three measures was used in analyses.

### fNIRS data acquisition

fNIRS data were collected using the NTS topography system (Gowerlabs Ltd., London, UK). This system uses two continuous wavelengths of near-infrared light (780 and 850 nm) to detect changes in HbO_2_ and HHb concentrations, using a sampling rate of 10 Hz (75).

Infants were assessed using a custom-built fNIRS headgear consisting of two arrays, including 14 sources and 12 detectors to create a total of 34 channels, covering bilateral frontal, inferior-frontal and temporal regions, with a source-detector separation of 2 cm (**Figure 1B** and **Figure 1C**). Infants underwent several fNIRS tasks as part of the BRIGHT project, focusing on social processing, deferred imitation, and habituation response.

Therefore, brain regions for data acquisition were chosen to maximize the likelihood of recording meaningful data for all tasks (76).

For each infant, head measurements were taken to enable the alignment of the headgear with the 10–20 placement coordinates (77). Over the front of the head, the headgear was placed so that a vertical red line marking the centre of the band aligned with the nasion with the silicon band laying just above, and in line with, the eyebrows. The headgear was placed so that the reference optode (the third lower optode from the back) was placed over the tragus.

To model for the headgear placement offline, photographs of the participants (one from the front and one from either side) were taken after the headgear was placed and again at the end of the fNIRS session. To assess the headgear placement, photographs of the participants had to be sufficiently good in quality (i.e. not blurred) and with the camera pointing at the ear, at a position directly perpendicular to the infant’s anatomical reference on the side of the head being photographed.

Infants were excluded from further analysis if the band was excessively high over the front above the eyebrows, by checking the displacement of the lower edge of the band with respect of the eyebrows with a “traffic-light” code (red, orange, green). Three experimenters independently classified the infants’ frontal headgear placement, and discrepancies were discussed until agreement was reached. Channels of those infants with valid placement over the front but horizontal lateral displacement between the ideal and the actual position of the reference (optode marked in green in **Figure 1B**) equal or greater than 1.6 cm were renumbered, so that each channel was shifted either backward or forward one full channel location in space

At each visit, the fNIRS data acquisition took place while infants sat on their parent’s lap. The parent was instructed to refrain from interacting with the infant during the stimuli presentation. Videos were of male and female Gambian adults singing nursery rhymes in Mandinka (for 70 seconds) and videos of toys in action (for 60 seconds), video blocks alternated, with each type shown to the infant twice, with the aim of keeping the infant awake, calm, attentive and as still as possible. The functional connectivity (FC) paradigm formed part of a greater battery of fNIRS paradigms that ran in a continuous and consistent order across all infants. The fNIRS FC acquisition lasted for a maximum of 520 seconds (two rounds of 260 seconds each) or were stopped as soon as the participant became fussy. Stimuli were presented using a MATLAB® custom-written stimulus presentation framework, Task Engine (sites.google.com/site/taskenginedoc) and Psychtoolbox on an Apple Macintosh computer.

### fNIRS preprocessing and data-analysis

Data analyses were carried out using in-house programmes developed in MATLAB (MathWorks, Natick, MA) (**Figure SI4** summarises fNIRS data preprocessing steps). Stringent multi-level data-quality assessments were used to ensure all data included in the group were of adequate quality. First, low-quality channels based on physiological indicators of quality were pruned using QT-NIRS (https://github.com/lpollonini/qt-nirs). The software takes advantage of the fact that the cardiac pulsation is also recorded in addition to haemodynamic activity associated with brain-driven variance including task-based responses and those contributing to functional connectivity (78), and quantifies its strength in the spectral and temporal domains with two measures: the Scalp Coupling Index (SCI) and the Peak Spectral Power (PSP) (for more details, see (79). For each infant, data quality was assessed channel-by-channel, and SCI and PSP were calculated with sliding 3-second windows (threshold SCI=0.70, threshold PSP=0.1, empirically defined and based on (80)) (79). Channels that had both SCI and PSP below threshold for more than 70% of the windows (79), were excluded from further pre-processing steps.

Raw intensity data were converted to optical density via a logarithm after dividing by the temporal mean of each channel (*hmrIntensity2OD*.*m* function from Homer2 tool (81)) and bandpass filtered (0.009 – 3 Hz) (*hmrBandpassFilt*.*m* function from Homer2 (81)). Using first a wide filter leaves in the signal contamination from the high bandwidth information (i.e. respiration, pulse, etc.) that we aim to regress out next. To help manage the systemic arterial signal that is well known to contaminate fNIRS signals (82), we regressed out the mean of the signals across the array, a method also known as global signal regression (GSR) (83, 84) (Results of analysis performed also without GSR are reported in the Supplementary Materials). Hereafter, optical density data underwent a second bandpass filtered (0.009 – 0.08 Hz), as previously described (47). Motion artefacts were then rejected using global variance of temporal derivatives (GVTD, STD=5) in the NeuroDOT toolbox (https://www.nitrc.org/projects/neurodot/) (85) (**Figure SI5** show how we tested the effect of different STD thresholds of GVTD on data inclusion). To further manage potential motion contamination of the data, we additionally removed five seconds before and after each motion artefact, and only chunks of data that were at least 20 seconds long were retained.

Optical density data were converted to relative concentrations of haemoglobin using the modified Beer–Lambert law (*hmrOD2Conc*.*m* function from Homer2 (81)). Differential path length factors were calculated (86) and adapted to wavelengths (780 and 850 nm) and ages (5 months = 5.24 and 4.25; 8 months = 5.26 and 4.26; 12 months = 5.27 and 4.28;18 months = 5.30 and 4.30; 24 months = 5.32 and 4.32).

To increase the statistical power of our analysis and to reduce the number of multiple comparisons for all statistical analysis to investigate developmental changes in connectivity, we averaged the concentration changes of the channels that survived pre-processing into sections. The sections were established via the 17 channels of each hemisphere which were grouped into front, middle and back (for a total of six regions) based on a previous co-registration of the BRIGHT fNIRS arrays onto age-appropriate templates (71) (**Figure SI1**). Each section represented a reasonable estimate of an anatomically consistent array coverage across the group, and was used to increase statistical power while minimizing errors due to variability in head anatomy and cap placement. For each participant, the Pearson-r correlation matrix between all the sections was calculated for both HbO_2_ and HHb, resulting in a 6×6 matrix of section-pair correlations. We then applied a Fisher z-transformation on the correlation matrix for further statistical analyses. Infants with at least 250 seconds of valid data after pre-processing were considered for further analyses. To choose the optimal minimum amount of valid data, per each infant at each time point for both the HbO_2_ and HHb signal, we correlated the connectivity matrices estimated in the first portion of data with the one in the last portion of data by increasing the seconds of data considered (i.e. the first 60 seconds with the last 60 seconds, the first 100 seconds with the last 100 seconds, the first 120 seconds with the last 120 seconds, etc.). We found that correlating the first and the last 250 seconds of valid data after pre-processing provided the highest percentage of infants with strong correlation between the first and the last portion of data, suggesting that FC patterns reached a usable stability after 250 seconds (**Figure SI6**).

### Cognitive flexibility at preschool age

Gambian infants from BRIGHT were cross-sectionally assessed at the age of 3 or 5 years for cognitive flexibility using the “card sorting” task from the tablet-based Early Years Toolbox (87) (http://www.eytoolbox.com.au). In this task, children were asked to sort cards (i.e. red rabbits or blue boats) either by colour (red or blue) or shape (rabbits or boats), and to flexibly switch back and forth between these rules (88).

Due to disruptions to field work as a result of a political crisis in December 2016 - January 2017 that forced us to pause recruitment, data were collected in two ranges of preschool ages, younger preschoolers (age mean ± SD=47.96 ± 2.77 months, N=77) and older preschoolers (age mean ± SD=57.58 ± 2.11 months, N=84) (76).

### Statistical Analyses

To investigate developmental trajectories of functional connectivity between 5 and 24 months, we ran linear mixed models (89) which is the standard statistical test to analyse repeated measures and longitudinal data with dropouts (90). Statistical analysis was performed with SPSS Version 28.0 statistic software package. Compared to repeated measures ANOVA, linear mixed models account for within-participant dependence and allow for missing data by using only information from the individual at the other visits (91). All possible interhemispheric homotopic, intrahemispheric within section, fronto-posterior, and crossed connections between the six regions were inserted as dependent variables in a linear mixed model, for a total of 21 linear mixed models. Functional connectivity was modelled as the dependent variable as a function of age with a random participant effect and random errors. Intercept and age were fixed effects, while within-participant dependence was modelled as a random effect (92). This same procedure was used in other longitudinal studies that explored developmental changes over time (31, 93, 94). The linear mixed model included the 132 infants who had valid data for at least two visits (55 infants contributed with data from two visits, 51 infants contributed with data from three visits, 22 infants contributed with data from four visits, four infants contributed with data from all of the five visits). The type of covariance between the observations was specified as Autoregression (AR) as two measures close in time of the same participant are likely to be correlated (95). To ensure statistical reliability, significant results from the linear mixed models were corrected for multiple comparisons using Bonferroni correction (p=0.00238).

Anthropometric measures were converted to age and sex adjusted z-scores that are based on World Health Organization Child Growth Standards (96). Weight-for-Length (WLZ) and Head Circumference (HCZ) z-scores were computed. Delta WLZ (ΔWLZ) were calculated for various periods (0-1 months, 0-5 months, etc.) by subtracting the z-score at the later age from that at the previous age. The use of delta z-scores as outcome measures allowed us to assess the impact of positive or negative deviation from the expected growth trajectories. To investigate the impact of undernutrition on FC development, we used ΔWLZ as independent variables in regression analyses on HbO_2_ (as the chromophore with the highest signal-to-noise ratio) FC at 24 months, our final time point of data collection. To maximise power, we considered only those FC that showed a statistically significant change with age. These analyses were adjusted by WLZ at birth and HCZ at 7/14 days, to more confidently assume that the associations between growth and FC were specific to the impact of change in WLZ postnatally and not confounded by the size or maturity of the infant at birth. We used HCZ measured at 7/14 days and not HCZ at birth as HC measures at birth are unreliable following vaginal delivery. Therefore, HCZ at 7/14 days is a more accurate head measure than HCAZ at birth. To ensure statistical reliability, results from the regression analyses on each FC were corrected for multiple comparisons using false discovery rate (FDR)(97) per each connection investigated, i.e. 21 possible ΔWLZ values per each connection.

To investigate whether FC early in life predicted cognitive flexibility at preschool age, we regressed later cognitive flexibility against FC that showed a significant change across the first two years of life.

## Acknowledgments

We thank the parents and infants who took part in this study without whom this work would not have been possible. We also thank the broader team of staff at the MRC Keneba Field Station for supporting us in the collection of this data.

## Funding

- This study was supported by: a Bill & Melinda Gates Foundation Grant OPP1127625; core funding MC-A760-5QX00 to the International Nutrition Group by the Medical Research Council UK; the UK Department for International Development (DfID) under the MRC/DfID Concordant agreement.
- CB is supported by a Leverhulme Trust Early Career Fellowship (ECF-2021-174).
- BM is supported by an ESRC Secondary Data Analysis Initiative Grant (ES/V016601/1).
- SEM and SMC are supported by a Wellcome Trust Senior Research Fellowship award to SEM (220225/Z/20/Z).
- AE is supported by funding from the National Institute for Mental Health (R01MH122751).

## Author contribution

Conceptualization: CB, SEM, SLF, CEE, ATE Methodology: CB, AB, SMC, BM, SEM, SLF, CEE, ATE

Investigation: SMC, GG, EM, ET, TF, LA Visualization: CB, ATE

Supervision: SEM, SLF, CEE, ATE Writing-original draft: CB, ATE

Writing-review & editing: SB, SMC, BM, SEM, SLF, CEE, ATE

## Competing interests

Chiara Bulgarelli is a research consultant for Gowerlabs Ltd., the company that produces the NTS optical topography system used in this work.

## Data and materials availability

Data supporting this paper will be made available subject to established data sharing agreements.

## Notes

### Summary of Updates

We added further clarifications on several core data analysis steps, such as coregistration and global signal regression. We also performed the additional analyses requested by the reviewers, exploring the associations between changes in connectivity and changes in growth and cognitive flexibility.

