## Supplementary material for "Growth in early infancy drives optimal brain functional connectivity which predicts cognitive flexibility in later childhood": SI

**This file includes:**

**Supplementary Results:**

Linear mixed model results showing the developmental trajectories of functional connectivity in Gambian infants over the first 2 years of life age (fNIRS pre-processing without global signal regression) 2

**Supplementary Figures:**

Figure SI1 – Schematic representation of the fNIRS array and the functional connections tested 3

Figure SI2 - Linear mixed models result showing FC that displayed a statistically significant change with age (fNIRS pre-processing without global signal regression) 4

Figure SI3 – Some examples of scatterplots showing the association between ΔWLZ and FC at 24 months 5

Figure SI4 – fNIRS preprocessing steps 6

Figure SI5 – The effect of different thresholds for GVTD for motion detecion and minium valid data after pre-processing on data inclusion 7

Figure SI 6 – Strength of correlation between FC in the first and last portion of data 8

**Supplementary Tables:**

Table SI1 - FC that significantly changed with age (fNIRS pre-processing without global signal regression) 9

Table SI2 - Results of the regression analyses of the effect of FC on cognitive flexibility in younger and older preschoolers 10

Table SI3 - Results of the correlational analyses between changes in growth (ΔWLZ) and cognitive flexibility in younger and older preschoolers 11

Table SI4. Results of the correlational analyses between changes in functional connectivity between 5 and 24 months and cognitive flexibility in younger and older preschoolers 12

Table SI5 - Characteristics of included and excluded participants and seconds of data included in the analyses at each age 13

### Supplementary Results

**Linear mixed model results showing the developmental trajectories of functional connectivity in Gambian infants over the first 2 years of life age (fNIRS pre-processing without global signal regression)**

To rule out that the global signal regression (GSR) performed as part of our fNIRS preprocessing might had an impact on our results, we re-processed the data without this step. All the other steps were kept the same. Hereafter, we reperformed the linear mixed models (LMM) on all the possible 21 interhemispheric homotopic, intrahemispheric within section, fronto-posterior, and crossed connections.

Results on the Fisher-z transformed correlation coefficients (z-RHO scores) on the oxygenated haemoglobin ($HbO_2$) showed that left ($F=10.4$, $p<0.001$) and right fronto-middle ($F=4.82$, $p<0.001$) FC increased with age. Results on the Fisher-z transformed correlation coefficients (z-RHO scores) on the deoxygenated haemoglobin (HHb) showed that showed that frontal interhemispheric FC decreased with age ($F=3.5$, $p<0.002$) (**Table SI1** and **Figure SI2**).

### Supplementary Figures

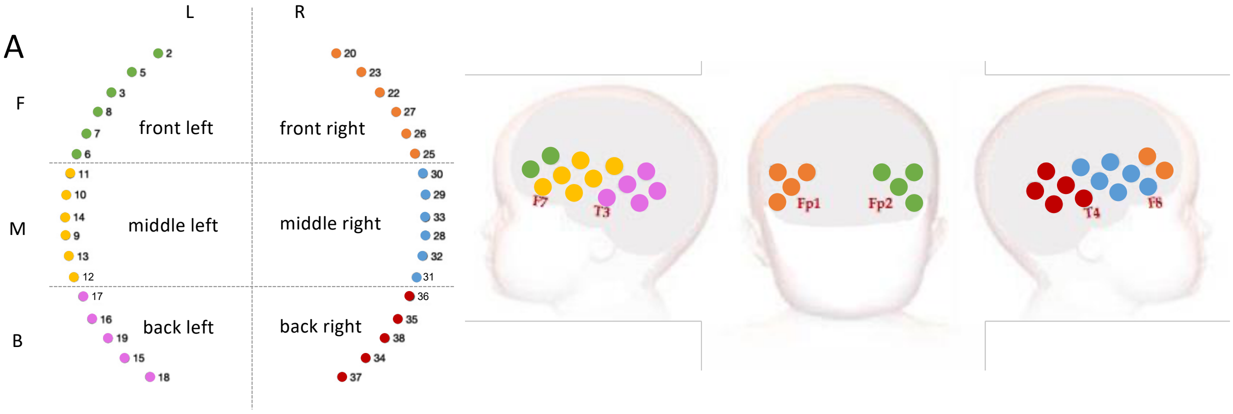


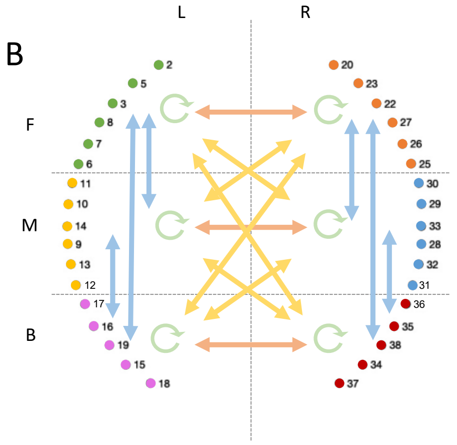


| | Left Front | Right Front | Middle Left | Right Middle | Left Back | Right Back |
|---|---|---|---|---|---|---|
| Left Front | Left Front Intra | Front Inter | Left Fronto-Middle | Cross Left Front – middle right | Left Fronto-Back | Cross Left Front – Right Back |
| Right Front | | Right Front Intra | Cross Right Front – Left Middle | Right Fronto-Middle | Cross Right Front – Left Back | Right Fronto-Back |
| Left Middle | | | Left Middle Intra | Middle Inter | Left Middle-Back | Cross Left Middle – Right Back |
| Right Middle | | | | Right Middle Intra | Cross Right Middle – Left Back | Right Middle-Back |
| Left Back | | | | | Left Back Intra | Back Inter |
| Right Back | | | | | | Right Back Intra |

**Figure SI1. Schematic representation of the fNIRS array and the functional connections tested**. (**A**) Each dot represents a channel, colours on the left plot corresponds to colours on the right on the baby's head. 6 sections. (**B**) The 21 connections tested in the linear mixed models. Interhemispheric homotopic connections are in orange (connecting the same regions between hemispheres, i.e., front left with front right), intrahemispheric connections within section are in green (correlations of channels belonging to the same region), fronto-posterior are in blue (connecting front and middle, middle and back, and front and back regions of the same hemisphere), and crossing interhemispheric connections (interhemispheric non-homotopic, connecting the front and middle, middle and back, and front and back regions of the two hemispheres) are in yellow.

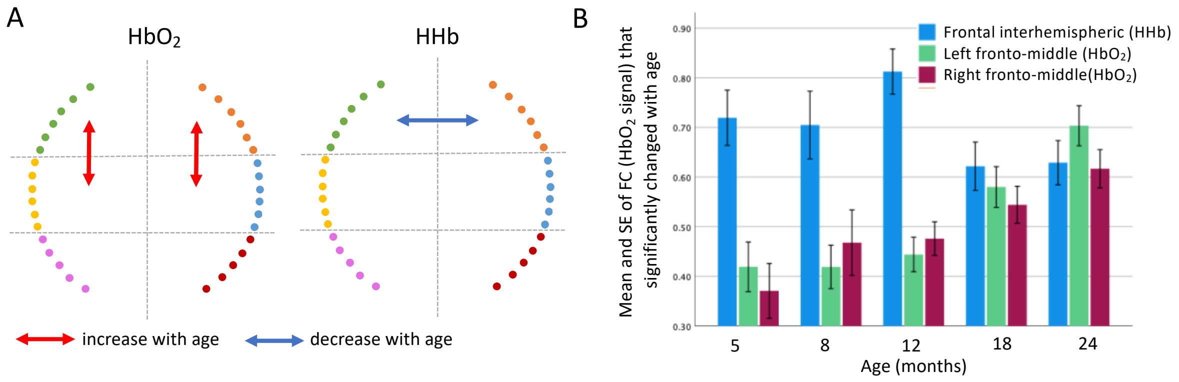


**Figure SI2. Linear mixed models result showing FC that displayed a statistically significant change with age (fNIRS pre-processing without global signal regression).** (**A**) Results of the linear mixed model, blue indicates connections that decreased with age, red indicates connections that increase with age. (**B**) Mean and SE of the functional connections that changed with age. Error bars are 1 SE.

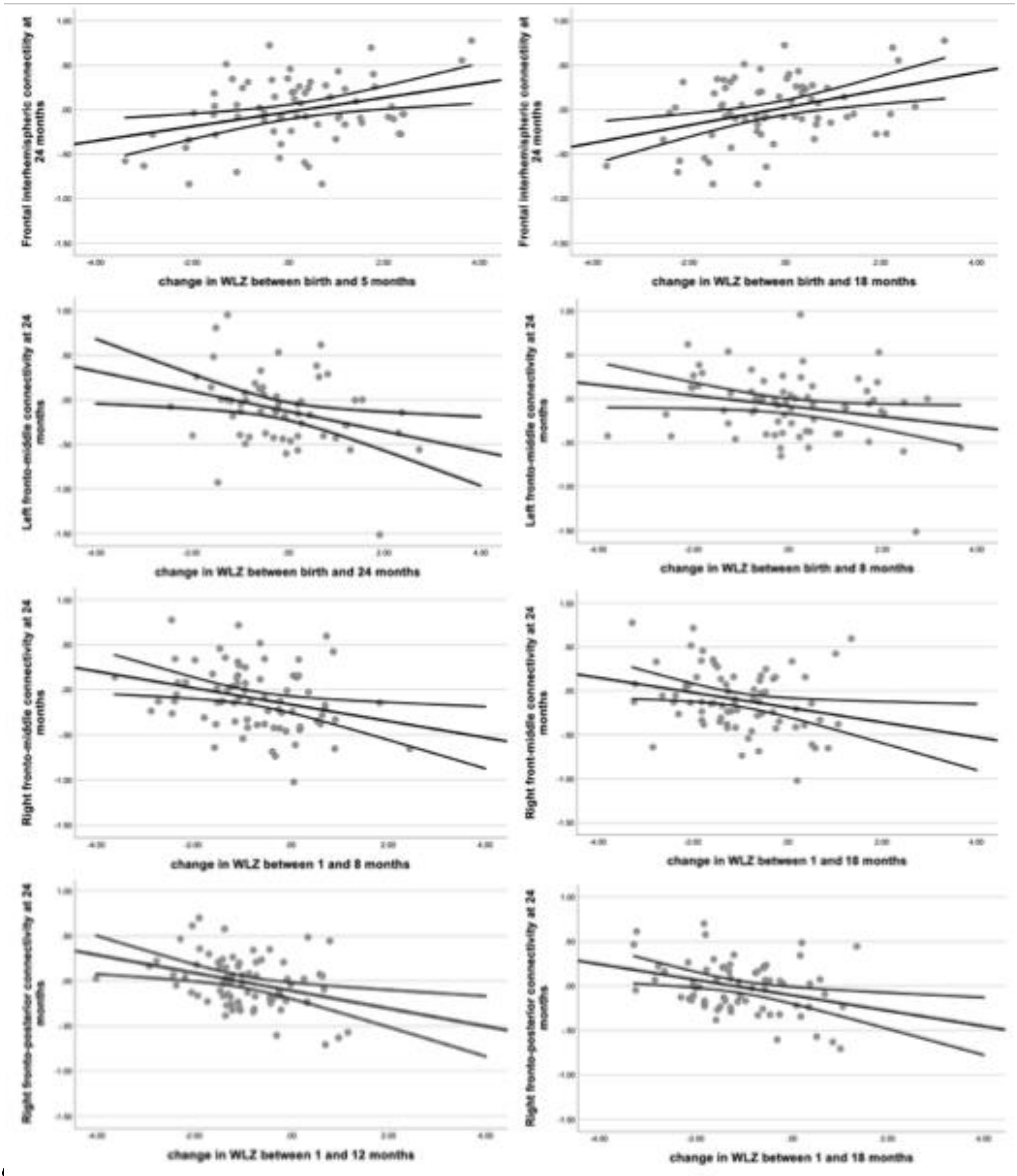


**Figure SI3. Some examples of scatterplots showing the association between ΔWLZ and FC at 24 months.** The black lines represent the line of best fit and the 95% confidence intervals.

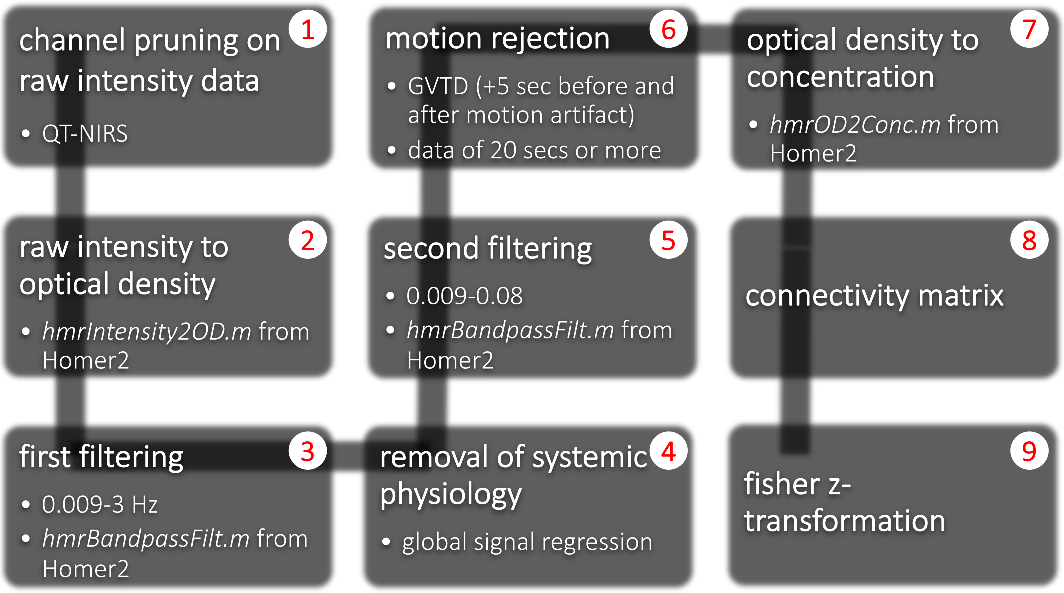


**Figure SI4. fNIRS preprocessing steps**. The column “infants included in the analyses” in Table 3 refers to those participants whose data survived these preprocessing steps.

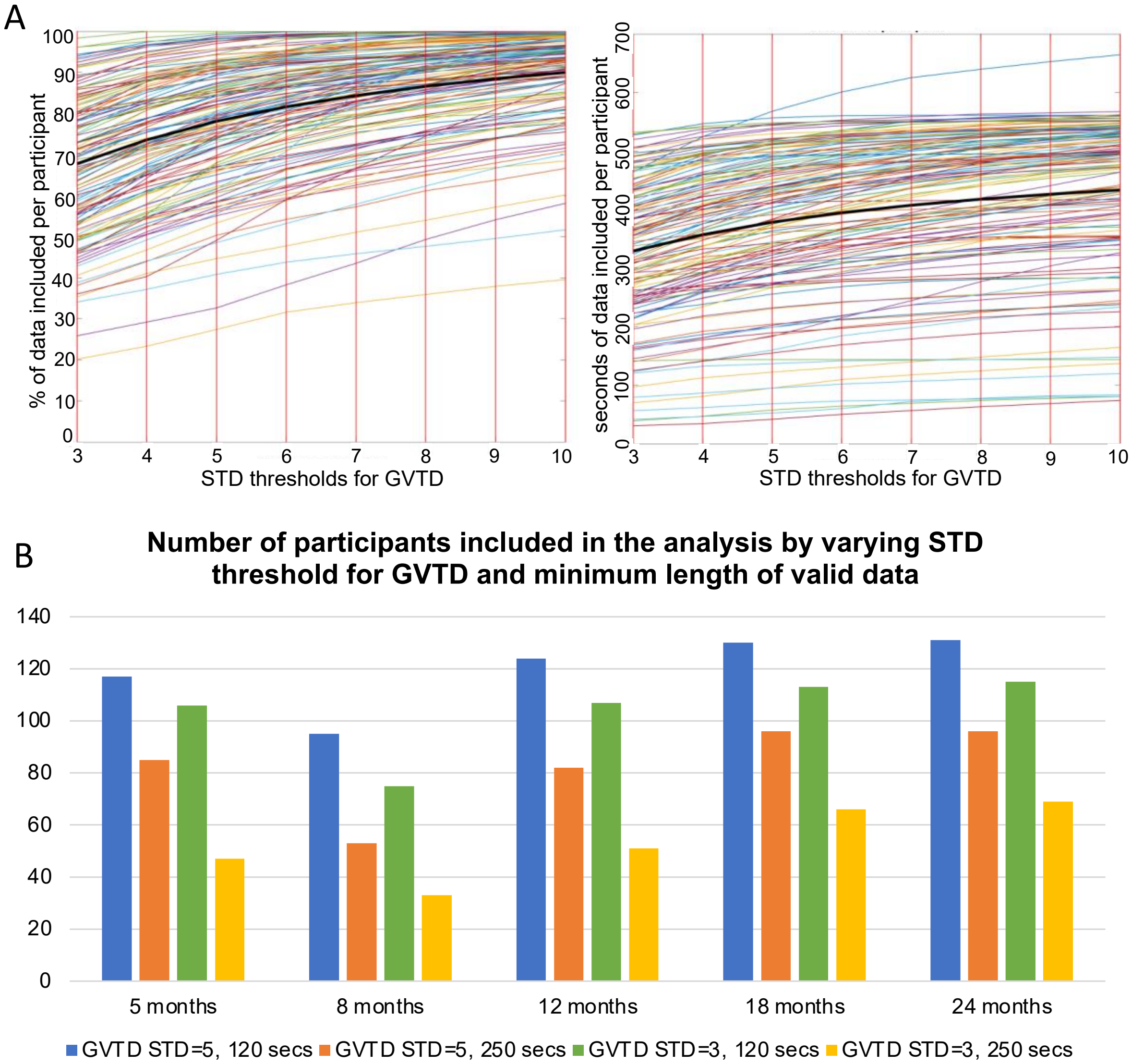


**Figure SI5**. **The effect of different thresholds for GVTD for motion detection and minimum valid data after pre-processing on data inclusion.** (A) Percentage of data included (left) and seconds of data included (right) per participant. Each line represents an infant, the black line represents the mean value. These graphs are reported from the 12 months sample as example. (B) Number of infants included in the LMM by varying STD threshold for GVTD and minimum length of valid data at different ages.

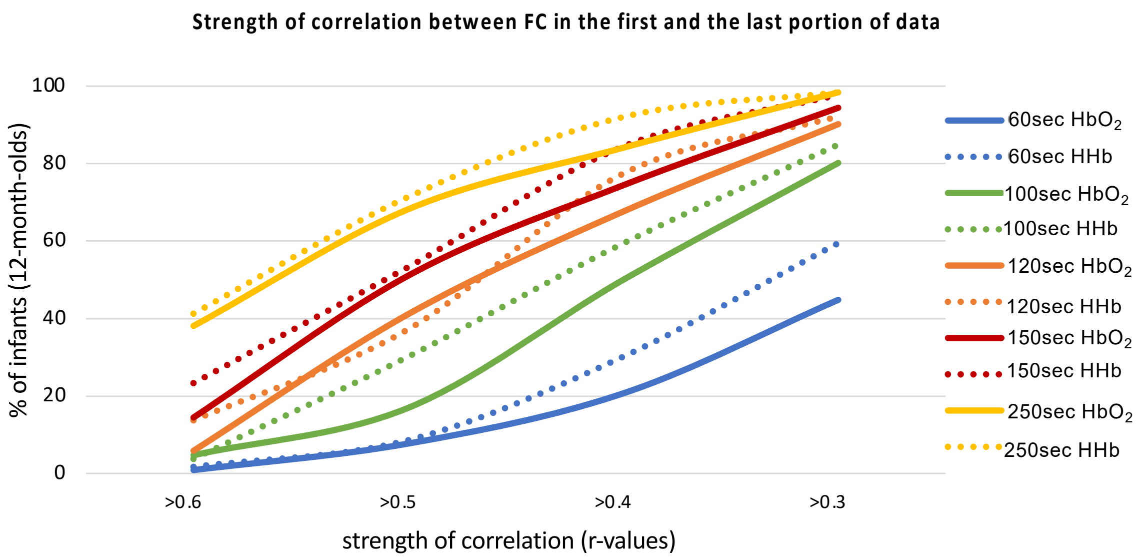


**Figure SI6**. **Strength of correlation between FC in the first and last portion of data**. This graph is reported from the age 12 month sample as example.

### Supplementary Tables

**Table SI1. FC that significantly changed with age (fNIRS pre-processing without global signal regression).** Results are displayed in terms of estimated betas, standard errors and p values.

| **FC** | ***F*** | ***p*** | **Baseline (5 months) Betas (SE), *p*** | **5-8 change Betas (SE), *p*** | **5-12 change Betas (SE), *p*** | **5-18 change Betas (SE), *p*** | **5-24 change Betas (SE), *p*** |
|---|---|---|---|---|---|---|---|
| **$HbO_2$** | | | | | | | |
| Left fronto-middle | 10.4 | <0.001 | -0.55 (0.29), <0.063 | 0.05 (0.05), 0.239 | 0.07 (0.04), 0.125 | 0.16 (0.04), <0.001 | 0.26 (0.04), <0.001 |
| Right fronto-middle | 4.82 | <0.001 | -0.24 (0.28), 0.398 | 0.03 (0.05), 0.535 | 0.05 (0.04), 0.246 | 0.08 (0.04), 0.051 | 0.18 (0.04), <0.001 |
| **HHb** | | | | | | | |
| Frontal interhemispheric | 3.5 | 0.002 | 0.18 (0.33) 0.591 | -0.07 (0.06), 0.260 | -0.07 (0.05), 0.214 | -0.17 (0.05), 0.002 | -0.17 (0.05), 0.002 |

**Table SI2**. **Results of the regression analyses of the effect of FC on cognitive flexibility in younger and older preschoolers.** Significant positive associations are in green, significant negative associations are in orange, and non-significant (NS) associations are in blue.

| | | **Cognitive flexibility in younger preschoolers** | **Cognitive flexibility in older preschoolers** |
|---|---|---|---|
| **Frontal interhemispheric FC** | **5 mo** | F(1,38)=4.21, p=0.047, R2=0.102 | NS |
| | **8 mo** | NS | NS |
| | **12 mo** | NS | NS |
| | **18 mo** | NS | F(1,45)=4.82, p=0.030, R2=0.115 |
| | **24 mo** | NS | NS |
| **Left fronto-middle FC** | **5 mo** | NS | NS |
| | **8 mo** | NS | NS |
| | **12 mo** | F(1,33)=7.86, p=0.009, R2=0.197 | NS |
| | **18 mo** | NS | F(1,45)=5.72, p=0.021, R2=0.115 |
| | **24 mo** | NS | NS |
| **Right fronto-middle FC** | **5 mo** | NS | NS |
| | **8 mo** | NS | NS |
| | **12 mo** | F(1,38)=4.82, p=0.034, R2=0.115 | NS |
| | **18 mo** | NS | NS |
| | **24 mo** | NS | NS |
| **Right frontal-posterior FC** | **5 mo** | NS | NS |
| | **8 mo** | F(1,32)=5.6, p=0.024, R2=0.153 | NS |
| | **12 mo** | NS | NS |
| | **18 mo** | NS | NS |
| | **24 mo** | NS | F(1,49)=4.85, p=0.032, R2=0.092 |

**Table SI3. Results of the correlational analyses between changes in growth (∆WLZ) and cognitive flexibility in younger and older preschoolers.**

| **Cognitive Flexibility in younger preschoolers** | | | | | | | |
|---|---|---|---|---|---|---|---|
| **∆WLZ** | **birth** | **1 mo** | **5 mo** | **8 mo** | **12 mo** | **18 mo** | **24 mo** |
| **birth** | | r(54)=-0.169, p=0.214 | r(51)=-0.167, p=0.232 | r(51)=-0.249, p=0.072 | r(53)=-0.198, p=0.147 | r(51)=-0.151, p=0.280 | r(41)=-0.241, p=0.102 |
| **1 mo** | | | r(64)=0.031, p=0.805 | r(63)=0.001, p=0.993 | r(66)=0.013, p=0.916 | r(62)=0.084, p=0.511 | r(48)=-0.021, p=0.885 |
| **5 mo** | | | | r(64)=-0.075, p=0.548 | r(66)=-0.032, p=0.798 | r(63)=0.008, p=0.947 | r(50)=-0.091, p=0.520 |
| **8 mo** | | | | | r(66)=-0.044, p=0.719 | r(62)=0.067, p=0.600 | r(50)=-0.041, p=0.775 |
| **12 mo** | | | | | | r(64)=0.097, p=0.439 | r(51)=-0.012, p=0.932 |
| **18 mo** | | | | | | | r(49)=-0.107, p=0.454 |
| **24 mo** | | | | | | | |

| **Cognitive flexibility in older preschoolers** | | | | | | | |
|---|---|---|---|---|---|---|---|
| **∆WLZ** | **birth** | **1 mo** | **5 mo** | **8 mo** | **12 mo** | **18 mo** | **24 mo** |
| **birth** | | r(55)=0.036, p=0.789 | r(58)=-0.122, p=0.353 | r(56)=-0.138, p=0.303 | r(56)=-0.153, p=0.252 | r(55)=-0.149, p=0.268 | r(53)=-0.137, p=0.319 |
| **1 mo** | | | r(64)=-0.138, p=0.268 | r(61)=-0.113, p=0.379 | r(62)=-0.113, p=0.294 | r(59)=-0.013, p=0.919 | r(59)=-0.069, p=0.597 |
| **5 mo** | | | | r(72)=-0.002, p=0.984 | r(73)=-0.025, p=0.829 | r(68)=0,064, p=0.601 | r(70)=0.046, p=0.704 |
| **8 mo** | | | | | r(72)=0.026, p=0.827 | r(68)=0.098, p=0.421 | r(69)=0.034, p=0.777 |
| **12 mo** | | | | | | r(68)=0.091, p=0.456 | r(70)=-0.057, p=0.634 |
| **18 mo** | | | | | | | r(65)=-0.103, p=0.406 |
| **24 mo** | | | | | | | |

**Table SI4. Results of the correlational analyses between changes in functional connectivity between 5 and 24 months and cognitive flexibility in younger and older preschoolers.**

| | **Cognitive Flexibility** | |
|---|---|---|
| **ΔFC between 5 and 24 mo** | **Younger preschoolers** | **Older preschoolers** |
| Frontal interhemispheric connectivity | r(12)=-0.368, p=0.196 | r(20)=-0.188, p=0.401 |
| Left fronto-middle FC | r(10)=0.253, p=0.428 | r(17)=0.270, p=0.264 |
| Right fronto-middle FC | r(12)=0.248, p=0.392 | r(20)=0.192, p=0.392 |
| Right frontal-posterior FC | r(12)=0.188, p=0.520 | r(20)=0.164, p=0.465 |

**Table SI5. Characteristics of included and excluded participants and seconds of data included in the analyses at each age**. WD=withdrawn, D=deceased, MV=missed visit, DD=developmental delay, NIRS not undertaken = the participant was assessed but did not want or could not perform the NIRS assessments, FC not undertaken = the participant was assessed with other NIRS task, but not FC, Fussed out = the participant wore the headband and the FC acquisition had started but the participant showed signs of fussiness soon after the start of the acquisition, MP=missing pictures of the headband placement, EM=missing event markers, TI=technical issues during the NIRS testing session. The proportion of children included in the analysis was computed based on the infants with FC data.

| Age | N - % | Not tested | | | | NIRS not undertaken | FC not undertaken | **Infants with FC data** | Fussed out | Experimental errors | | | Headband Placement | Too many channels excluded | Not enough data after pre-processing | **Infants included in the analyses** | **Seconds of data included in the analyses (mean±SD)** | **Inclusion rate (from the 204 infants recruited)** |
|---|---|---|---|---|---|---|---|---|---|---|---|---|---|---|---|---|---|---|
| | | WD | D | MV | DD | | | | | MP | EM | TI | | | | | | |
| **5 months** | N | 2 | 1 | 2 | 3 | 10 | 7 | **179** | 16 | 11 | 3 | 3 | 7 | 5 | 47 | **87** | 382.68±92.78 | 42% |
| | % | 0.98 | 0.49 | 0.98 | 1.47 | 4.90 | 3.43 | **87.74** | 8.93 | 5.39 | 1.67 | 1.67 | 5.58 | 2.79 | 26.25 | **48.6** | | |
| **8 months** | N | 7 | 1 | 5 | 3 | 18 | 14 | **156** | 7 | 6 | 0 | 1 | 23 | 6 | 60 | **53** | 372.93±80.66 | 25% |
| | % | 3.43 | 0.49 | 2.45 | 1.47 | 8.82 | 6.86 | **76.47** | 4.48 | 3.84 | 0 | 0.64 | 14.7 | 3.84 | 38.46 | **33.97** | | |
| **12 months** | N | 9 | 1 | 4 | 3 | 17 | 13 | **157** | 4 | 1 | 0 | 2 | 10 | 2 | 56 | **82** | 372.93±80.66 | 40% |
| | % | 4.41 | 0.49 | 1.96 | 1.47 | 8.33 | 6.37 | **76.96** | 2.54 | 0.63 | 0 | 1.27 | 6.36 | 1.27 | 35.66 | **52.22** | | |
| **18 months** | N | 8 | 1 | 15 | 3 | 12 | 5 | **160** | 4 | 0 | 0 | 1 | 16 | 4 | 38 | **97** | 388.85±81.63 | 47% |
| | % | 3.92 | 0.49 | 7.35 | 1.47 | 5.88 | 2.45 | **78.43** | 2.5 | 0 | 0 | 0.62 | 10 | 2.5 | 23.75 | **60.62** | | |
| **24 months** | N | 13 | 1 | 29 | 1 | 3 | 4 | **153** | 0 | 2 | 0 | 0 | 6 | 4 | 45 | **96** | 399.56±79.86 | 47% |
| | % | 6.37 | 0.49 | 14.2 | 0.49 | 1.47 | 1.96 | **75** | 0 | 1.30 | 0 | 0 | 3.92 | 2.61 | 29.41 | **62.74** | | |
